# Effect of high-salt diet on mean arterial pressure, renal epithelial sodium channels and aquaporin subunits expression levels in Spontaneously Hypertensive Rats

**DOI:** 10.1101/630491

**Authors:** Chitra Devi Ramachandran, Khadijeh Gholami, Sau-Kuen Lam, Mohd Rais Mustafa, See-Ziau Hoe

## Abstract

An increase in blood pressure (BP) by a high-salt (HS) diet may involve the changes in the expression of epithelium sodium channels (ENaCs) and aquaporins (AQPs) in the kidney which affect the sodium- and water-handling mechanisms. In the present study, spontaneously hypertensive rats (SHRs) and Wistar Kyoto (WKY) rats were exposed to HS and regular-salt (RS) diets for 6 weeks and fluid intake was monitored. After 6 weeks, mean arterial pressure (MAP) and plasma hormonal activity of atrial natriuretic peptide (ANP), levels of angiotensin II (Ang II), aldosterone and arginine vasopressin (AVP) were determined. The expression of mRNA and protein levels of ENaC and AQP subunits in kidneys were quantified by real-time PCR and Western blotting. High-salt diet caused higher MAP only in SHRs and higher fluid intake in both strains of rats when compared with their respective controls on RS diet. The plasma levels of Ang II and aldosterone were low in both SHRs and WKY rats fed with HS diet. Meanwhile, plasma ANP activity was high in both strains of rats on HS diet; whilst the AVP showed vice versa effects. The renal expression of mRNA and protein levels of α- and γ-ENaCs was lowered by HS diet in both SHRs and WKY rats. Although β-ENaC mRNA and protein expression levels were depressed in SHRs but they were enhanced in WKY rats. On the other hand, AQP-1, 2 and 7 mRNA and protein expression levels were lowered in both strains of rats fed with HS diet, while that of AQP-3, 4 and 6 showed no significant changes. The suppression of mRNA and protein expression levels of ENaC and AQP subunits suggests that the HS-induced increase in the MAP of SHRs may not be due to the renal sodium and water retention solely.

## Introduction

Dietary salt (i.e. sodium chloride, NaCl) intake is the most remarkably modifiable environmental risk factor that attracts many studies on hypertension (HPN). It has been acknowledged as an important contributing factor of the aetiology and progression of HPN [1]. Despite the abundant experimental, interventional and epidemiological observations demonstrating an association between dietary salt and HPN, scepticism remains as to how high salt (HS) intake can be mechanistically linked to the increase in blood pressure (BP). Knowing the heterogeneity of HPN, it is likely to involve the intricate integration of multiple regulatory systems and the kidneys have long been implicated to play a central role in regulating BP. Defects in the kidneys sodium- and water-handling mechanisms have been mooted as one of the primary causes of HPN in HS intake [2].

The kidneys have the capacity to return altered BP to baseline level by increasing or decreasing sodium and water excretion in response to elevated or reduced BP [3]. This is accomplished in the kidney by the presence of renal membrane-bound protein i.e. epithelial sodium channel (ENaC) that fine-tune sodium reabsorption [4] and aquaporins (AQPs) that facilitate the transport of water and in some cases, other small uncharged solutes [5, 6].

Epithelial sodium channels (ENaCs) are composed of three homologous subunits i.e. the α, β and γ [7]. The α subunit is absolutely required for channel activity in that it is critical for the formation of ion the permeating pore, whereas β and γ subunits are necessary for maximal channel expression and activity at the cell surface and may also play a regulatory role [8]. Nevertheless, all the three subunits have significant effects on multimeric ENaC protein sodium transport capacity. The ENaC subunits are regulated by a variety of hormones especially aldosterone [9-11]. The aldosterone acts through mineralocorticoid receptor which in turn regulates ENaCs transcription [12, 13]. Beside aldosterone, arginine vasopressin (AVP), the major antidiuretic hormone (ADH), also acts as an antinatriuretic hormone that increases sodium reabsorption [14-16]. Apart from these two hormones, angiotensin II [17-19] has also been implicated with sodium transport. In addition, atrial natriuretic peptide (ANP) has been reported to be an inhibitor of ENaC [20]. Malfunctions of ENaC subunits affect their responses to dietary salt and thus, disturb sodium homeostasis. The functional role of ENaC in the development of salt-sensitive HPN (SSH) have been widely studied and a variety of responses have been reported [21-24]. Thus, investigation on ENaC and its role in sodium handling in response to HS diet intake are continually expanding.

Apart from sodium balance, the kidneys are also essential to maintain body water balance which also affects BP; and this is accomplished by the presence of aquaporins (AQP). Aquaporin (AQP) is a specialised transporter that allows cells to absorb a large amount of water needed to control the volume of both extra- and intracellular fluid. It was first discovered by Peter Agre in 1992 [25] and to date, 13 types of AQP subunits (AQP0 to AQP12) have been identified in mammals. The AQPs are found in different forms in the kidney i.e. AQP1, AQP2, AQP3, AQP4, AQP6 and AQP7 [26-29]. Renal AQPs are necessary for osmotic equilibration [30] and numerous studies been documented of the association between increased AQPs levels and pathogenesis of HPN [31, 32]. A physiologically relevant role in water reabsorption has been demonstrated for AQP1 to AQP4. Majority of the water absorption in the kidney occurs via AQP1, localised in the proximal tubule; and AQP2, expressed in the apical membrane of collecting duct [30, 33-35]. Similar to ENaCs, the expressions of AQPs in the kidneys were also found to be influenced by hormones.

In all the reported studies, inappropriate sodium and water retention by ENaC and AQP subunits have been shown to be involved in the pathogenesis of HPN. Most of the studies on the effect of HS were performed in Dahl salt-sensitive and salt-resistance rats as well as SD rats. But studies on the ENaCs and AQPs dysregulations in SHRs, the rat model that shares similar pathophysiology with essential HPN in human population, as a consequence of HS diet were far from complete. Therefore, in the present study, we used SHRs to investigate the expression level of both ENaC and AQP subunits as a result of HS intake. We hypothesised that chronic HS diet intake affects expressions of ENaC and AQP subunits in the kidney which lead to sodium and water retention, respectively, and the subsequent increase in BP.

## Materials and Methods

### Ethical approval

The study was carried out in the Department of Physiology and Medical Biotechnology Laboratory of the Faculty of Medicine, University of Malaya. All the experimental protocols involving animals and housing thereof were reviewed and approved by the Institutional Animal Care and Use Committee (IACUC) of the University of Malaya (Reference: 2014-01-07/Physio/R/HSZ) which maintains a full Association for Assessment and Accreditation of Laboratory Animal Care (AAALAC) accreditation.

### Experimental design and diet treatment

Male WKY rats and SHRs used in this study were bred at the University of Malaya Animal Experimental Unit from stock obtained from BioLASCO (Taiwan). After being weaned at 5 weeks of age, rats were housed in groups of 4 to 5 under controlled laboratory conditions (temperature 23 ± 5°C, 12:12 hour light/dark cycle and humidity 50% to 60%) with food and water provided *ad libitum* for at least 1 week prior to the onset of experimentation. Six-week-old WKY rats and SHRs were randomly assigned to receive food with either a regular salt (RS) content (0.2% w/v NaCl) or a high-salt (HS) content (4% w/v NaCl; Harlan Teklad, Germany) with free access of water. The potassium content in both diets was 0.6% (w/v). Four groups were thus studied:

Group 1: WKY receiving RS (WRS)

Group 2: WKY receiving HS (WHS)

Group 3: SHR receiving RS (SRS)

Group 4: SHR receiving HS (SHS)

The treatment period continued for 6 weeks. The water was added and replaced on alternate days.

### Measurement of mean arterial pressure (MAP) and fluid intake

Eight rats at the age of 12 weeks from each group were anesthetized with sodium pentobarbital (60mg/ kg; i.p.). The reflexes of the rat were checked, and it was placed on the rodent surgical table. A small incision (1.5 to 2 cm) was made in the neck for tracheostomy and carotid artery cannulation. The carotid artery was cannulated with a cannula pre-filled with heparinized normal saline (5IU/ ml) which was connected to a pressure transducer (MLT0380, ADInstrument). The transducer output was amplified as well as recorded continuously by Powerlab Data Acquisition System (ADInstrument, Sydney, Australia). The whole setup was allowed to stabilize for 30 to 45 minutes with the baseline recording carried out for 10 to 15 minutes. On the BP tracing, the up and down stroke waves represent the systolic and diastolic blood pressures, respectively. The mean arterial pressure (MAP) was also determined by using the formula of MAP = 1/3 (Systolic BP-Diastolic BP) + Diastolic BP. On the other hand, weekly intake of drinking fluids was estimated throughout the experimental period. The fluid intake was measured by subtracting the measured amounts provided to the remaining amounts in the cage.

### Plasma analysis

The plasma was obtained from blood samples collected from trunk blood in a chilled, peptidase inhibitor (for ANP) and heparinised (for Ang II, aldosterone and AVP) coated vacutainers by centrifugation at 3,000 rpm, 4°C for 20 minutes. Plasma ANP activity was quantified using radioimmunoassay (RIA) procedure as previously described by Gutkowska et al [36]. Data was expressed as pg/ml and the sensitivities of RIA and intra-as well as interassay coefficients of variation for ANP was 0.7pg/ml, 4.8% and 10%, respectively. Meanwhile, plasma Ang II (catalogue number: E-EL-R1430), aldosterone (catalogue number: ADI-900-173) and AVP (catalogue number: ADI-900-017A) levels were quantified by using a competitive enzyme-linked immunosorbent assay (ELISA) kits (Elabscience, China and Enzo Life Sciences, USA). All assays were performed according to the manufacturers’ guidelines with the lowest assay sensitivity limit of approximately 3.9pg/ml. Absorbance values were read at 405nm for aldosterone and 450nm for Ang II and AVP, using a microplate reader (Infinite M1000 Pro, Tecan, Switzerland).

### Tissue collection

At the end of the diet treatment i.e. at week 12, rats were euthanised (between the hours of 0800 to 1100) by a blow to the head, whole kidneys were harvested and snap frozen in dry ice. All tissues collected were stored at −80°C until further use.

### mRNA extraction, cDNA synthesis and quantitative reverse transcription polymerase chain reaction (qRT-PCR)

Kidneys weighing around 60mg were disrupted using rotor-stator homogeniser (Heidolph DIAX 600, Ballerup, Denmark) in Qiazol lysis buffer. Phase separation was initiated by adding chloroform (EMPARTA MERCK, Mumbai) and centrifuged at 12,000g for 15 minutes at 4°C. One volume of 70% (v/v) ethanol was added to upper aqueous phase and applied to RNeasy mini columns (Qiagen RNeasy Mini kit, Qiagen) and the remaining purification steps were carried according to manufacturer’s guidelines. Total RNA was quantified using Nanodrop (Thermo Scientific NanoDrop 2000). A total 500ng of RNA was reverse transcribed into cDNA by using Bio-Rad iScript Reverse Transcription Supermix for RT-qPCR (Biorad, Hercules, CA, USA) according to manufacturer’s instruction. The steady-state of ENaCs and AQPs expression level in kidney’s mRNA was measured using RT-qPCR. All primers for α-ENaC encoded by *Scnn1a* (5′-CCTAAGCCCAAGGGAGTTGA-3′ and 5′-ACACTACAAGGCTTCCGACA-3′), β-ENaC encoded by *Scnn1b* (5′-TGGACATTGGTCAGGAGGAC-3′ and 5′-AGCAGCACCCCAATAGAAGT-3′), γ-ENaC encoded by *Scnn1g* (5′-TGAGGCTTCCGAGAAATGGT-3′ and 5′-AATACTGTTGGCTGGGCTCT-3′), *AQP1* (5′-ACCCACTGGAGAGAAACCAG-3′ and 5′-AGAGTAGCGATGCTCAGACC-3′), *AQP2* (5′-AACTACCTGCTGTTCCCCTC-3′ and 5′-ACTTCACGTTCCTCCCAGTC-3′), *AQP3* (5′-GAACCCTGCTGTGACCTTTG-3′ and 5′-AGTGTGTAGATGGGCAGCTT-3′), *AQP4* (5′-ACACGAAAGATCAGCATCGC-3′ and 5′-TGACCAGGTAGAGGATCCCA-3′), *AQP6* (5′-GGATCTTCTGGGTAGGACCG-3′ and 5′-ACGGTCTTGGTGTCAGGAAA-3′), *AQP7* (5′-TATCTTCGCCATCACGGACA-3′ and 5′-CCCAAGAACGCAAACAAGGA-3′) and Gapdh (5′-GCTACACTGAGGACCAGGTT-3′ and 5′-TCATTGAGAGCAATGCCAGC-3′) were designed from NCBI official website (http://www.ncbi.nlm.nih.gov). All primers for target and endogenous control genes were obtained from Integrated DNA Technologies. The qRT-PCR reactions were carried out in triplicate in 96-well plates and each PCR sample consisted of 6μl 2X SYBR green master mix buffer (Roche), 0.024μl of forward and reverse primers 25nmole and 3.953μl of RNase-free water. The reactions were performed using the Applied Biosystems StepOnePlus Real-Time PCR System and fold change (FC) was assessed by establishing a delta-delta cycle threshold (Ct) between Gapdh, the calibrator gene and target genes. The average Ct values of target and calibrator genes obtained from qRT-PCR instrumentation were imported into a Microsoft Excel spreadsheet and the ΔΔCt was calculated using the equation Ct _Target_ – Ct _Gapdh_ as described by Livak *et al*. [37].

### Protein extraction, quantification and immunoblotting

Frozen kidneys tissue weighing approximately 80mg was cut into small pieces which were then submerged in 800ml radioimmunoprecipitation assay (RIPA) buffer solution (BioVision, Country) containing protease and phosphatase inhibitors at a ratio of 1:10. The mixture was homogenised for 30 seconds using rotor-stator homogeniser (Heidolph DIAX 600, Ballerup, Denmark). The total protein of the kidneys was extracted by centrifugation at 14,000g for 15 minutes at 4°C and protein concentration was determined using micro bicinchoninic acid (BCA) protein assay kit (Thermo Scientific, Rockford, Il, USA) according to manufacturer’s guidelines. An equal amount of protein was separated with 8% (v/v) and 12% (v/v) sodium dodecyl sulphate polyacrylamide gel electrophoresis (SDS-PAGE) for ENaC subunits and AQPs, respectively and transferred onto polyvinylidene fluoride (PVDF) membrane (BioRad, USA). Upon blocking the membrane with 2% (w/v) Amersham ECL Prime Blocking Reagent (GE Healthcare) for an hour at room temperature, the membranes were then probed with primary antibodies (AB3530P, SC25354, AB3534P for α-, β- and γ-ENaCs, respectively; AB3272, AB3066, AB3276, AB3594, AB3073 and AB15568 for AQP1, 2, 3, 4, 6 and 7, respectively and ABS16 for Gapdh) diluted in 0.1% (v/v) Tween20/PBS (PBST) at 4°C overnight. This followed with incubation in appropriate secondary antibodies conjugated with horseradish peroxidase (HRP) (Abcam, USA) for an hour at room temperature. The blots were then developed using Super Signal West Pico Chemiluminescent Substrate (Thermo Scientific, Rockford, Il, USA) and signals were captured by using high sensitive CCD camera-based imager (BioSpectrum Imaging System). The band intensity of each target was analysed using Image J software and protein expression level was expressed as a ratio to Gapdh (loading control). All experiments were carried out in triplicate and average band intensities were then determined.

### Statistical analysis

Statistical analysis was performed using GraphPad Prism (GraphPad Software, La Jolla, CA, USA). All data are expressed as the mean ± standard error of means (SEM) of 4 to 8 rats. Comparisons between groups (SHS vs SRS, and WHS vs WRS) were performs by independent unpaired Student’s *t*-test. The differences were considered statistically significant at *p* values <0.05.

## Results

### Effect of high-salt (HS) diet on mean arterial pressure (MAP) and fluid intake

As shown in Fig 1, SHRs consuming HS diet (SHS) developed a significantly (p<0.001) higher MAP (188.44 ± 4.66mmHg) as compared with SHRs consuming RS diet (SRS) which displayed a MAP of 163.20 ± 4.72mmHg. The MAP of WKY rats, on the other hand, did not show any significant difference between HS and RS groups.

**Fig 1:**
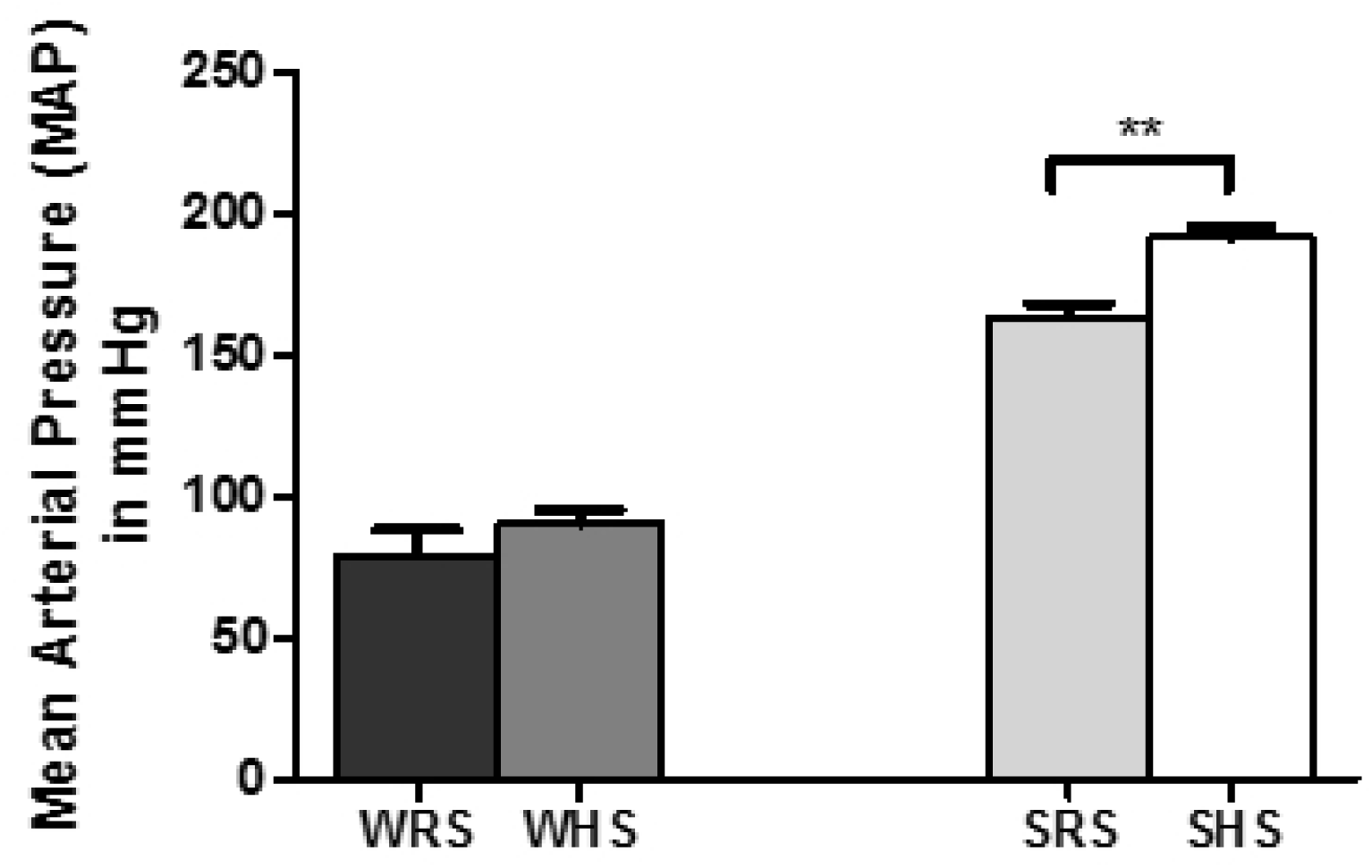
Effect of HS diet on mean arterial pressure (MAP) in SHRs and WKY rats. Data was presented as mean ± SEM; n = 8 rats. The **p<0.01 SHS compared with SRS using Student’s *t*-test. Abbreviation: WRS: WKY rats fed with RS; WHS: WKY rats fed with HS; SRS: SHRs fed with RS; SHS: SHRs fed with HS.

Fig 2 shows higher fluid intake by both SHS and WHS when compared with their relevant control groups, SRS and WRS, respectively. The water intake in SHS was 279.06 ± 39.22ml equated with SRS that drank 138.21 ± 6.17ml (p<0.01). Meanwhile, WHS consumed about 296.44 ± 24.70ml when compared with 149.23 ± 18.41ml by WRS (p<0.001).

**Fig 2:**
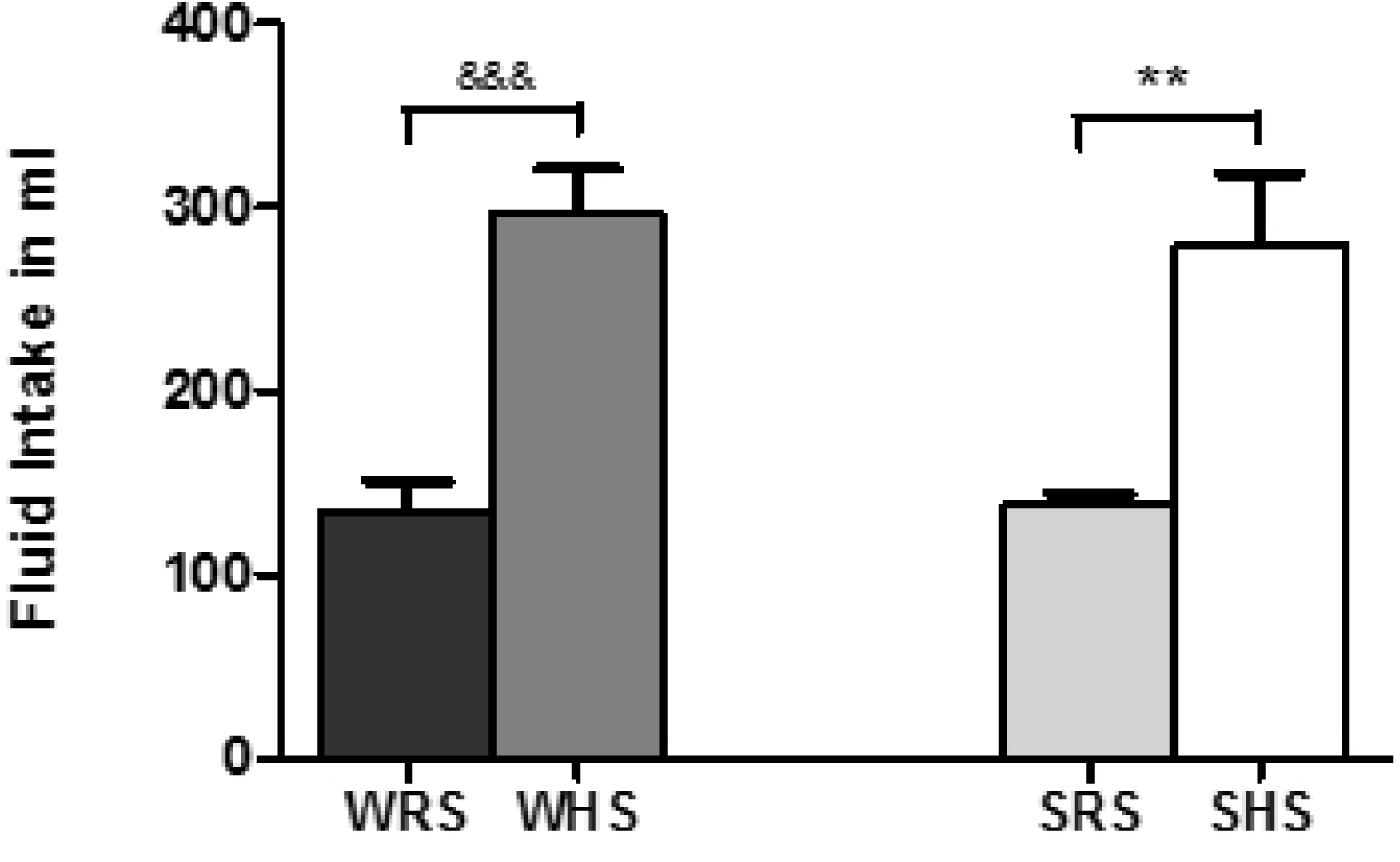
Effect of HS diet on fluid intake in SHRs and WKY rats. Data was presented as mean ± SEM; n = 8 rats. The **p<0.01 SHS compared with SRS; ^&&&^p<0.001 WHS compared with WRS using Student’s *t*-test. Abbreviation: WRS: WKY rats fed with RS; WHS: WKY rats fed with HS; SRS: SHRs fed with RS; SHS: SHRs fed with HS.

### Plasma analysis

The plasma ANP activity of SHS, on the other hand, was significantly augmented compared to SRS (p<0.01). The plasma ANP activity in SHS was 80.73 ± 10.14pg/ml whilst 46.39 ± 7.06pg/ml in SRS, an increase of nearly 50% of the activity level. In addition, WHS (41.53 ± 5.81pg/ml) also showed a significant higher plasma ANP activity relative to WRS (p<0.001) with a great fold change (Fig 3A).

**Fig 3:**
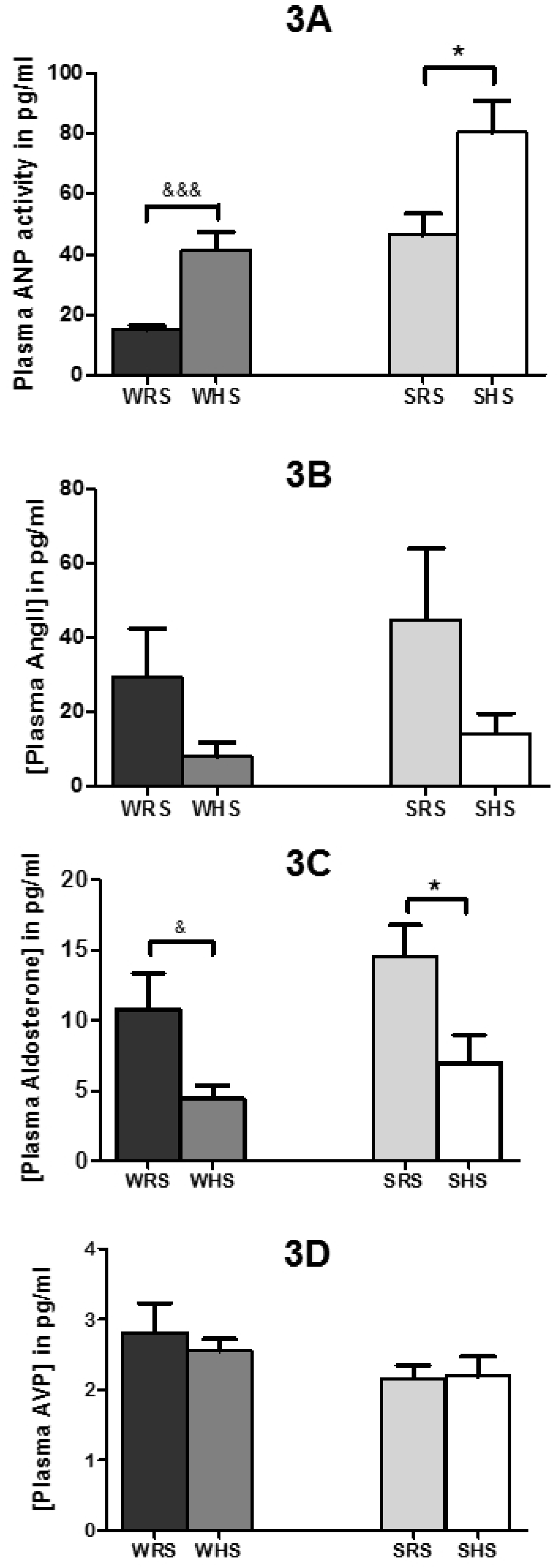
Effect of HS diet on plasma (A) atrial natriuretic peptide (ANP) activity, (B) angiotensin II, (C) aldosterone and (D) AVP levels in SHRs and WKY rats. Data presented as mean ± SEM; n = 6 rats. The *p<0.05 SHS compared with SRS, ^&^p<0.05 and ^&&&^p<0.001 WHS compared with WRS using Student’s *t*-test. Abbreviations: WRS: WKY rats fed with RS; WHS: WKY rats fed with HS; SRS: SHRs fed with RS; SHS: SHRs fed with HS

Meanwhile, as shown in Fig 3B, the plasma Ang II level in SHRs and WKY rats fed with HS diet (SHS and WHS) were lower when compared with SHRs and WKY rats on RS diet (SRS and WRS), respectively. As expected, both SHRs and WKY rats fed with the HS diet showed lower plasma aldosterone level when compared with their respective control groups. The plasma aldosterone level of SHS was 7.00 ± 1.92pg/ml when compared with SRS with 14.55 ± 2.25pg/ml (p<0.05); whilst in WHS the plasma aldosterone level was 5.27 ± 1.16pg/ml when compared with 10.80 ± 3.27pg/ml in WRS (p<0.05) (Fig 3C).

As shown in Fig 3D, the plasma AVP of SHS was only slightly higher compared with SRS whilst it was lower in WHS to WRS. However, the results were not significant.

### Effect of HS diet on mRNA expression levels of ENaC subunits in the kidney

The HS diet was found to be able to lower the mRNA expression levels of *Scnn1a* gene encoding α-ENaC in the kidneys of both SHRs and WKY rats when compared between their counterparts i.e. SHS vs SRS and WHS vs WRS (p<0.01), respectively, as evidenced in Fig 4A. Meanwhile, *Scnn1g*, gene encoding γ-ENaC, was found to be significantly (p<0.01) lower in WHS when compared with WRS. The expression of γ-ENaC was also downregulated in SHS when compared with SRS; however, the result was not significant (Fig 4C). On the other hand, the mRNA expression level of β-ENaC was found to be also lower in SHS when compared with SRS; however, the results were not significant. Meanwhile, there was no significant change in the expression of β-ENaC in WHS when compared with WRS (Fig 4B).

**Fig 4:**
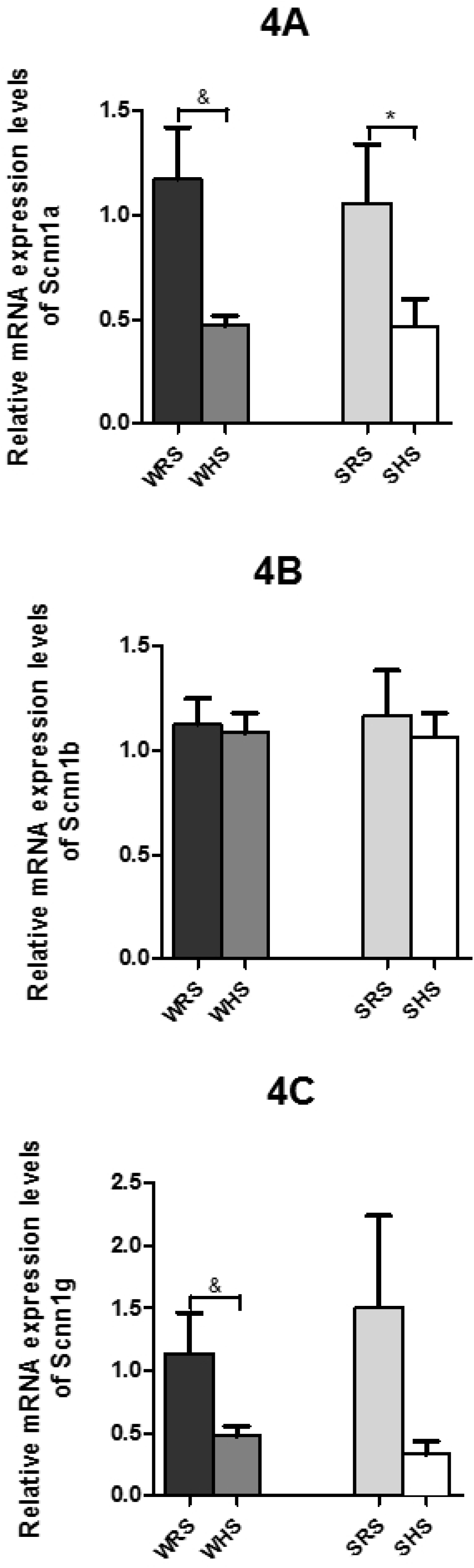
Relative mRNA expression levels of (A) *Scnn1a* encoding α-ENaC, (B) *Scnn1b* encoding β-ENaC and (C) *Scnn1g* encoding γ-ENaC in the kidneys under the influence of HS diet. Data are presented as mean ± SEM; n = 4 rats. The *p<0.05 SHS compared with SRS and ^&^p<0.05 WHS compared with WRS using Student’s *t*-test. Abbreviations: WRS: WKY rats fed with RS; WHS: WKY rats fed with HS; SRS: SHRs fed with RS; SHS: SHRs fed with HS.

### Effect of HS diet on mRNA expression levels of AQP in the kidney

The expression levels of *AQP1* (Fig 5A) and *AQP7* (Fig 5F) were markedly lower with p<0.05 and p<0.01, respectively in WKY rats being fed with HS diet when compared with WKY rats on RS diet. Meanwhile, the SHRs did not show significant expression change of parallel comparison. Meanwhile, the level of *AQP2* was also found to be significantly lower in SHS when compared with SRS (p<0.05) (Fig 5B). However, no significant differences in the expression level of other AQPs, i.e. *AQP3*, *AQP4* and *AQP6* between different groups of animals were observed (Fig 5C, D, and E).

**Fig 5:**
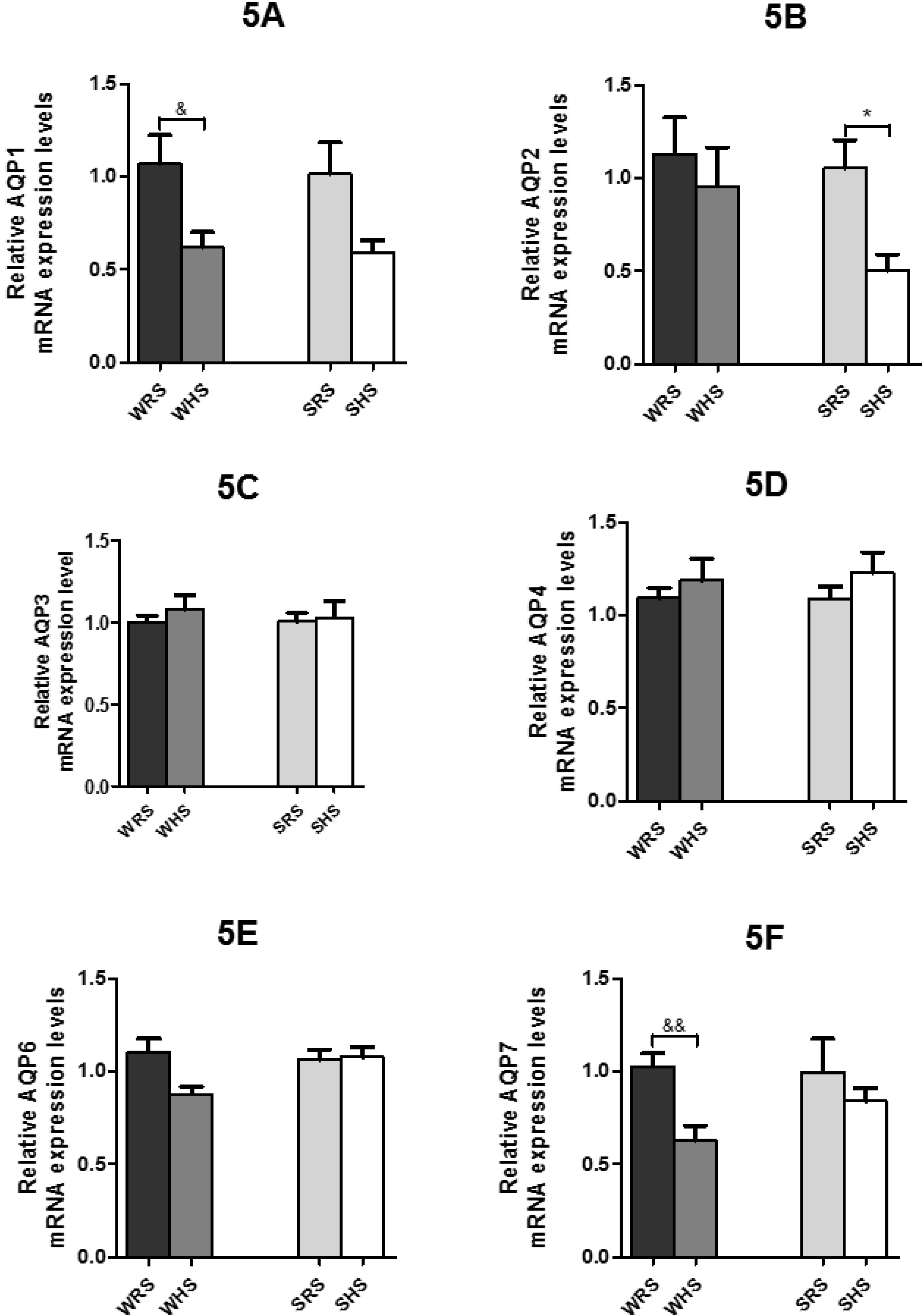
Relative mRNA expression levels of (A) *AQP1*, (B) *AQP2*, (C) *AQP3*, (D) *AQP4*, (E) *AQP6* and (F) *AQP7* in the kidneys under the influence of HS diet. Data are presented as mean ± SEM; n = 4 rats. The *p<0.05 SHS compared with SRS, ^&^p<0.05 and ^&&^p<0.01 WHS compared with WRS using Student’s *t*-test. Abbreviations: WRS: WKY rats fed with RS; WHS: WKY rats fed with HS; SRS: SHRs fed with RS; SHS: SHRs fed with HS.

### Effect of HS diet on protein expression levels of ENaC subunits in the kidney

esults in Fig 6 shows that HS diet depressed the protein expression level of α-, β- and γ-ENaC in SHRs. However, significant (p<0.05) depression was only observed in the γ-ENaC (Fig 6C). In the WKY rats, on the other hand, HS diet caused suppressions α- and γ-ENaCs expression but enhancement of β-ENaC expression. However, all the expressed changes in WKY rats were not significant.

**Fig 6:**
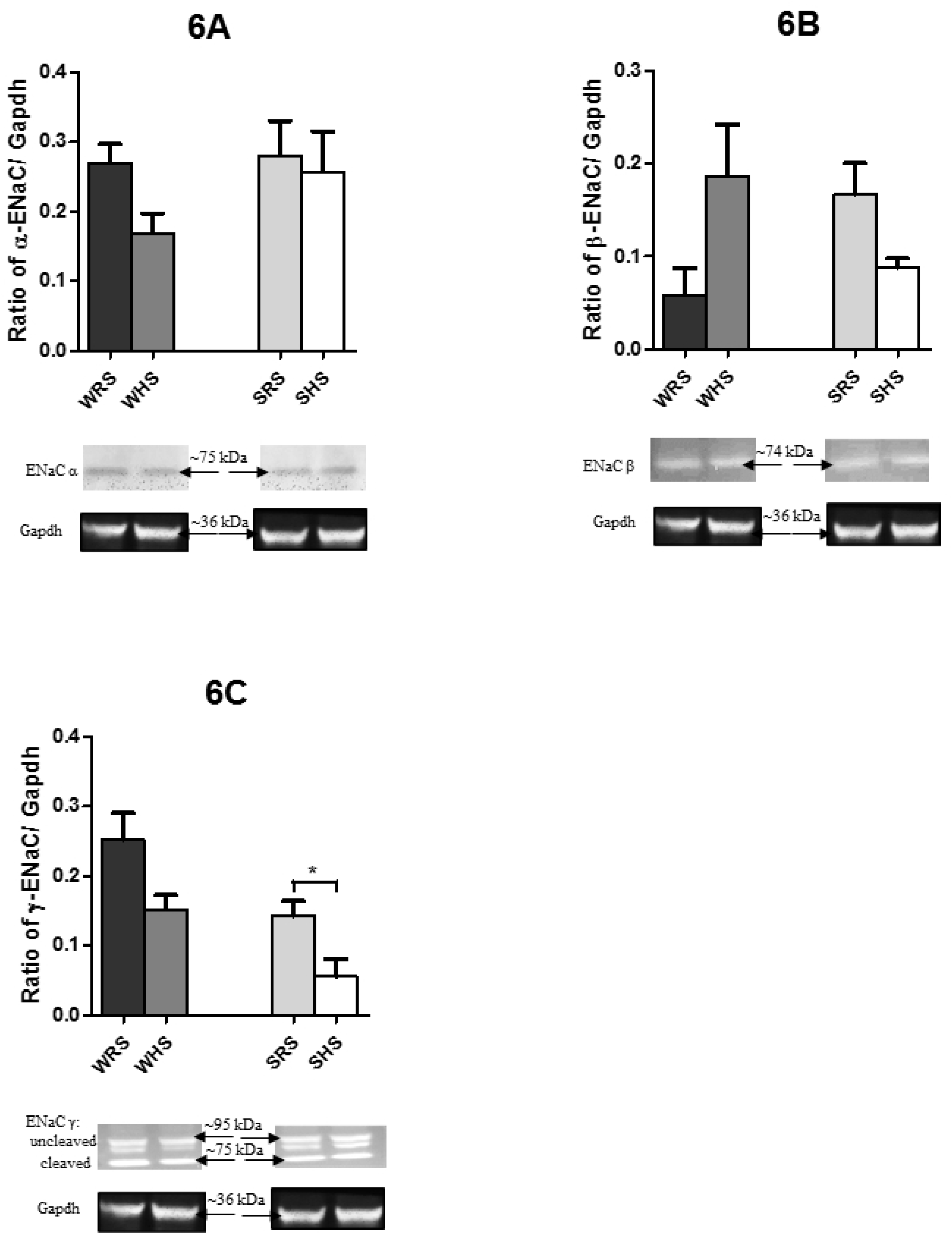
Protein expression levels of (A) α-ENaC, (B) β-ENaC and (C) γ-ENaC in the kidneys under the influence of HS diet. Data are presented as mean ± SEM; n = 4 rats. The *p<0.05 SHS compared with SRS using Student’s *t*-test. Abbreviations: WRS: WKY rats fed with RS; WHS: WKY rats fed with HS; SRS: SHRs fed with RS; SHS: SHRs fed with HS.

### Effect of HS diet on protein expression levels of AQP in the kidney

As shown in Fig 7A to F, HS diet was found to lower the protein expression levels of AQP1, AQP2 and AQP7 in both SHRs and WKY rats compared with their counterparts. In contrast, the AQP3 expression level was enhanced in both strains of rats. On the other hand, the AQP4 and AQP6 protein expression levels were contra-expressed in SHRs and WKY rats i.e. the protein expression level was enhanced in WKY rats but depressed in SHRs of being fed with HS diet.

**Fig 7:**
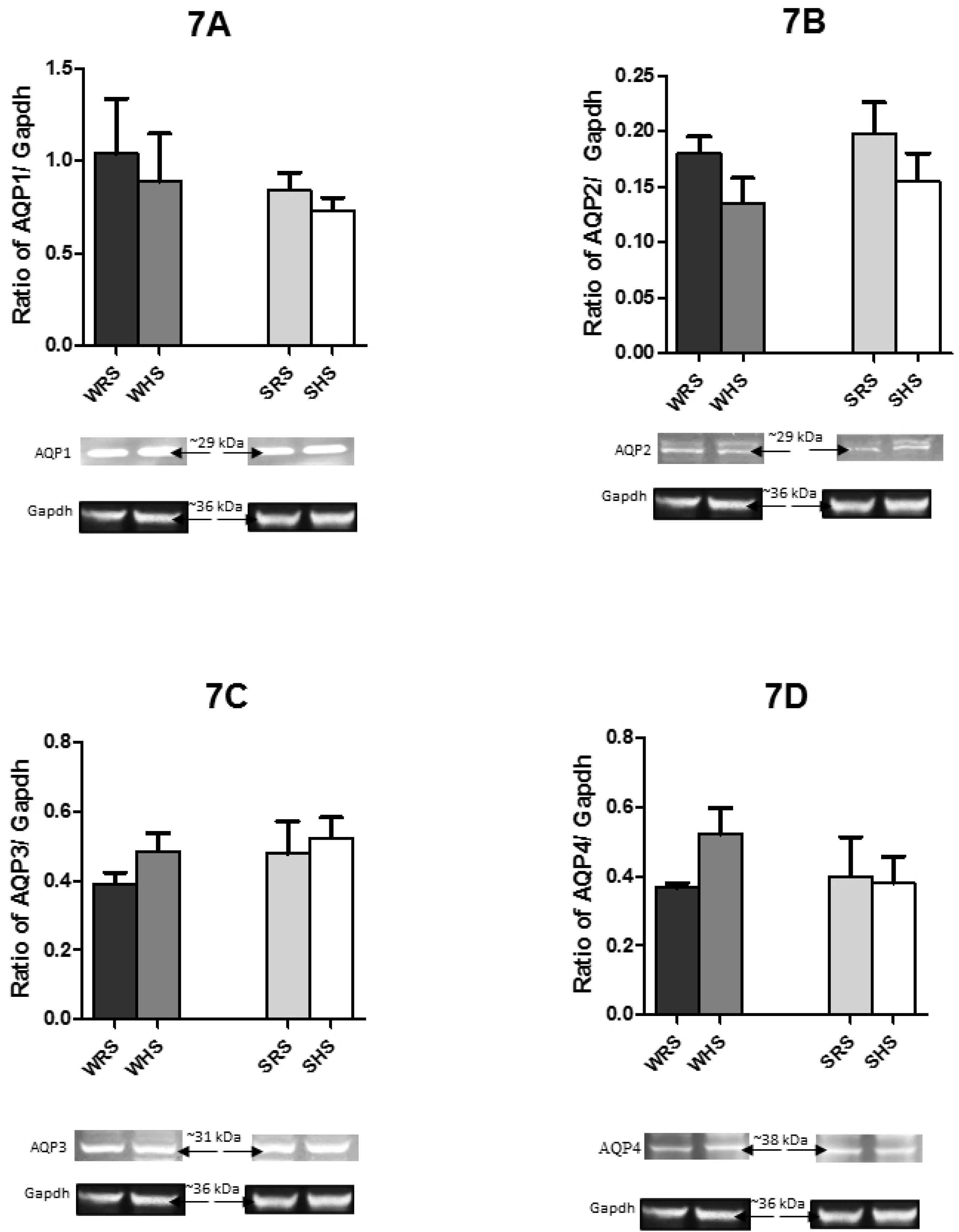
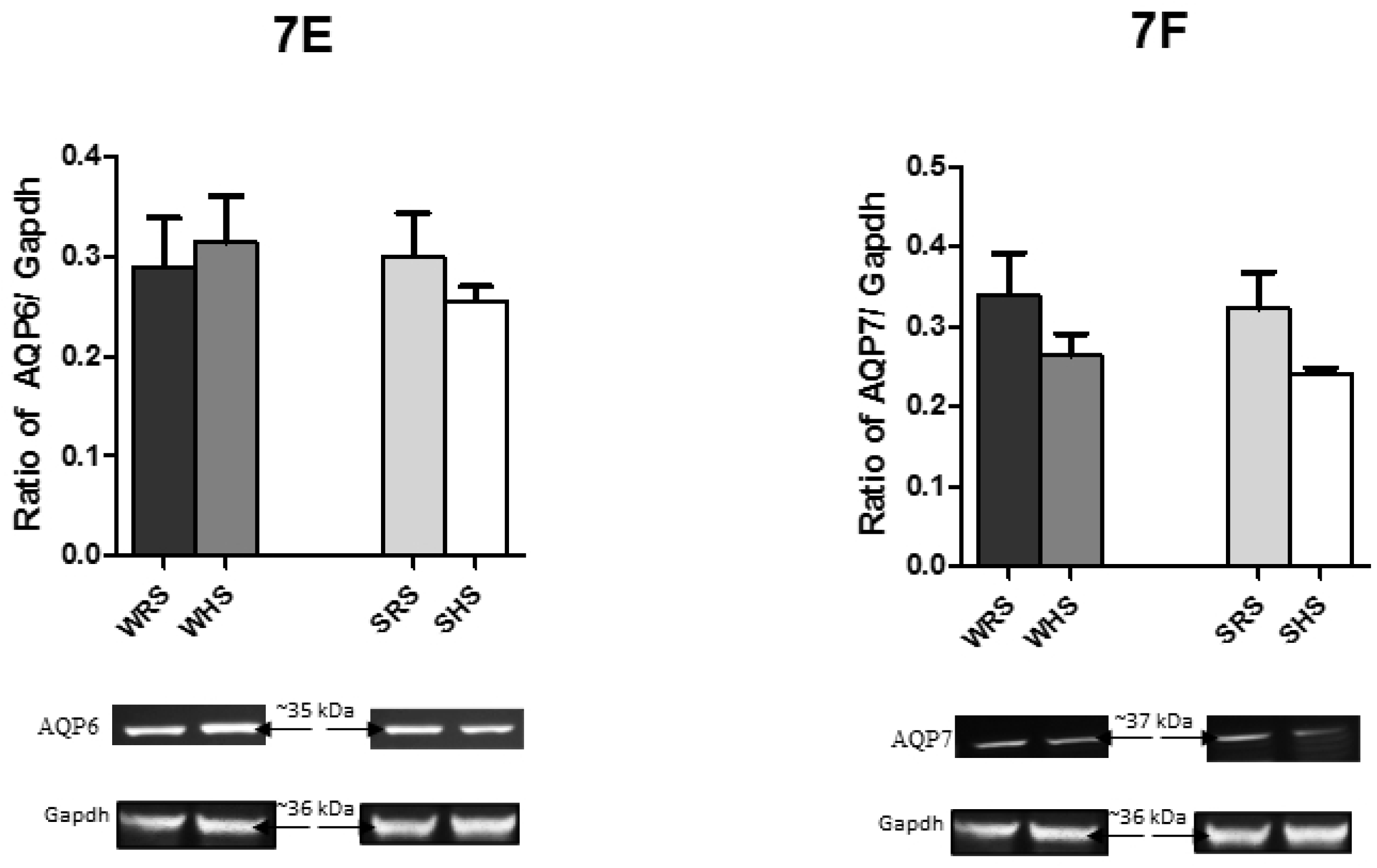
Protein expression levels of (A) AQP1, (B) AQP2, (C) AQP3, (D) AQP4, (E) AQP6 and (F) AQP7 in the kidneys under the influence of HS diet. Data are presented as mean ± SEM; n = 4 rats. Abbreviations: WRS: WKY rats fed with RS; WHS: WKY rats fed with HS; SRS: SHRs fed with RS; SHS: SHRs fed with HS.

## Discussion

In this present study, we found that SHRs on a HS diet showed significantly higher MAP when compared with SHRs on a RS diet (Fig 1). However, there was no significant difference between the MAP of WKY rats consuming of HS and RS diets, respectively. The current result is in accordance to the claim of SHRs to become hypertensive with normal/ RS intake as they have vascular smooth muscle cells that take up sodium excessively due to alteration of Na^+^-K^+^ pump [38]. Consequently, the intracellular sodium concentration ([Na^+^]_i_) increases and subsequently induces a rise in calcium concentration via sodium-calcium exchanger that further causes vasoconstriction. Therefore, an augmentation in sodium load such as high dietary salt intake is predicted to elevate the [Na^+^]_i_ even more [39], thus rises the MAP.

We also found that both the rat strains fed with HS showed higher fluid intake when compared to their respective controls (Fig 2). It is well known that the rise in plasma sodium content will increase plasma osmolarity, which causes a rise in extracellular fluid volume by promoting the transfer of fluid from intracellular to extracellular space as well as by stimulating the thirst centre [40, 41]. Thus, the elevation in the MAP of SHS in the present study may as a result of volume expansion under the influence of raised salt intake as reported by Qi et al. [42]. Therefore, the balance between salt and water in extracellular fluid is vital to ensure the precise regulation of osmolarity and thus the volume of body fluids, which in turn maintains the BP.

In the meantime, the present study showed higher plasma ANP activity in both SHRs and WKY rats fed with HS diet when compared with their respective controls (Fig 3A). It is well acknowledged that ANP is synthesised by atria in heart and secreted into the bloodstream in response to stretching of right atrial muscle cells by increased blood volume. In the bloodstream, the ANP act on distal convoluted tubule of nephron to inhibit sodium reabsorption and causes natriuresis [43, 44]. However, study by Greenwood et al. [45] showed a low circulating ANP in salt-loaded and water deprived Sprague-Dawley (SD) rats as compared with their euhydrated controls which contradict with the current result. Nevertheless, the present finding is in accordance with results from studies conducted by both Sagnella et al. [46] and Kohno et al. [47] which demonstrated higher plasma ANP levels during HS intake in patients with essential HPN and salt-sensitive patients, respectively. As ANP is an important indicator of blood volume; thus, the increase in plasma ANP in the present study may corroborate with our finding (Fig 2) that showed high fluid consumption of SHRs and WKY rats being fed with HS diet. A high fluid consumption due to HS intake would have increase ECF volume and this would have risen the stretch of cardiac chambers thus surge the secretion of ANP. Teleologically, the response of ANP would be logical in being protective against excessive sodium and water retention.

As expected, the plasma Ang II level in both strains of rat on HS diet was lower when compared to their respective controls (Fig 3B). Angiotensin II has been reported for its direct involvement in the control of renal sodium excretion and in neural control of sodium appetite, thus regulate body’s sodium balance [41, 48-51]. Our finding is in accordance to the findings of Greenwood et al. [45] that also showed decreased plasma Ang II concentration in salt-loaded SD rats; meanwhile, Mecawi et al [51] demonstrated an increased plasma Ang II in WKY rats fed with low-sodium diet. In the meantime, the plasma aldosterone level in both SHRs and WKY rats was lower when compared with their respective controls (Fig 3C). It is well documented that the synthesis of aldosterone increases in response to low plasma sodium so that sodium will be retained in the cell [52]. As such, the increase in sodium as in the present study would certainly secrete low aldosterone. Therefore, it is not surprising to see the low plasma aldosterone level in both SHRs and WKY rats fed with HS diet. The higher plasma aldosterone level in SHRs compared with WKY rats also explains the higher BP in SHRs as that of WKY rats. Furthermore, the release of aldosterone is also dependent on the level of Ang II; thus, the low Ang II might also lead to a low aldosterone level in both strains of rat.

Meanwhile, the plasma AVP level was found to be slightly higher in SHRs fed with HS salt diet whilst lower in WKY rats (Fig 3D). Generally, AVP causes vasoconstriction by acting on V_1_ receptor as well as promotes water reabsorption in the kidney via acting on a V_2_ receptor. The regulation of AVP is mainly by changes in osmolarity. Though AVP had been associated with the development and maintenance of salt-dependent and malignant forms of HPN as well as to influence baroreceptor reflexes, results regarding plasma AVP levels in hypertensive patients are found to be not consistent with high levels in some studies [53, 54] but normal or low levels in others [55]. Moreover, it has also been evidenced that enhanced thirst appeared to normalise plasma AVP concentrations in subjects on HS intake [53]. This may serve as the possible explanation in the present results as SHRs and WKY rats fed with HS diet showed higher fluid consumption compared to WKY rats of RS diet.

The effect of HS diet on the mRNA expression and protein distribution of ENaC subunits were also investigated in the present study. Both the mRNA and protein expression of α-, β- and γ-ENaC subunits were downregulated and lowered, respectively, in SHRs fed with HS diet when compared with SHRs on RS diet (Fig 4). The SHRs on normal or RS (0.2% Na^+^ content) diet has been reported to be able to retain excessive amount of sodium resulting from reduced glomerular filtration [56-58], enhanced tubular reabsorption [57, 59] and increased protein abundance of ENaC subunits in various part of kidney segments [57]. This in turn, contributes to the elevated BP in these rats. However, the 4% HS diet in the present study did not enhanced either the mRNA or protein level in SHRs suggesting that the HS diet induce compensatory natriuresis to maintain sodium homeostasis [21, 60] in SHRs. One of the compensatory natriuretic mechanism could be the low plasma Ang II as well as aldosterone levels and reduced in these to plasma proteins has been reported to lower α-ENaC mRNA level [60]. Aldosterone either from adrenal medulla stimulated by Ang II or from RAS, is known for its essential role in transcription of gene encoding α-ENaC, thus activating its activity [61, 62]. Therefore, our findings well correlate with reduced plasma Ang II and aldosterone levels with the low mRNA expression of the α-ENaC subunit and thus the lower protein content of α-ENaC.

Meanwhile, the mRNA expressions of β- and γ-ENaCs, which are known to be expressed independent of aldosterone [63, 64], were also found to be depressed in SHRs of being fed with HS diet. Activities of both β- and γ-ENaCs have been reported to be regulated by α-ENaC [65]. Hence, the low expression of β- and γ-ENaCs could be due to low level of α-ENaC. Therefore, it is postulated that co-expression of all ENaC subunits would result in a fully operating channel as their co-existence was required for maximal ENaC channel function [66]. This claim is further supported by the studies on the gene-knockout animal model in which the α-, β- and γ-ENaCs-knockout mice displayed metabolic abnormalities and death because of their lack of ability to retain sodium and water, as well as to excrete potassium [67]. Furthermore, the low plasma aldosterone and high plasma ANP of SHRs fed with HS diet (Fig 3A and 3C) may indicate that the high MAP in SHR caused by HS diet (Fig 1) was not due to alteration in the activity of ENaC and may involve other mechanisms such as activation of sympathetic nervous activity [4, 68, 69], enhancement of reactive oxygen species (ROS) [21, 70] and stimulation of cardiovascular control centre in brain. Subsequent to the low mRNA expression the protein content of β- and γ-ENaCs were also low in SHRs (Fig 6). On the other hand, the lower mRNA and protein levels of α- and γ-ENaCs in WKY rats fed with HS diet is in accordance to the claim that under physiological conditions, in normotensive rats (Dahl-salt-resistance/SD/WKY rats) there is neither no change in expression nor decreased expression of ENaC in the kidney in response to HS diet [22, 71-74]. However, the higher expression of β-ENaC protein level in the present study somehow needs further exploration.

In the present study, the mRNA of AQP showed various expression patterns in SHRs and WKY rats in response to HS diet. In SHS (SHRs being fed with HS diet), the mRNA levels of *AQP1*, *AQP2* and *AQP7* were found to be lower when compared to SRS (SHRs being fed with RS diet) (Fig 5). Consistent with the downregulation of *AQP1*, *AQP2* and *AQP7*, the protein levels of these AQPs were also depressed when compared with their controls (Fig 7). Similar changes in the mRNA and protein expressions of AQPs were demonstrated by WKY rats. Meanwhile, the mRNA expression levels of *AQP3* and *AQP4* (Fig 5C and D) were found to be enhanced in both strains of rats being fed with HS diet. However, the protein level of AQP4 (Fig 7D) in SHS was found to be lower when compared with its counterpart. Interestingly, AQP6 displayed contra-expression in mRNA and protein level in SHRs and WKY rats; in which mRNA level of SHR (SHS vs SRS) was enhanced but the protein level was depressed whilst mRNA level in WKY rats (WHS vs WRS) was lower but protein level was enhanced. All these observations in mRNA and protein levels of AQPs have led to interesting point to discuss.

Aquaporin1 is the major water channel in renal proximal tubule and loop of Henle that is responsible for reabsorbing 80% of glomerular filtrate [75, 76]. It has been reported that renal and cardiac AQP1 expressions were downregulated in conditions such as renal fibrosis in mice [77] and HS-induced HPN [78]. The present result is in accordance with the finding by Penna et al. [79] who showed that 8% HS downregulated APQ1. Hence, the downregulation of AQP1 in both SHRs and WKY rats could be interpreted as a compensatory mechanism to prevent larger water reabsorption in the proximal tubule and the consequent expansion of extracellular fluid volume [27]. It has been claimed that in SHRs, the AQP1 expression in kidney [32, 75] and brain [75, 80] to be upregulated. However, the HS diet in the present study showed downregulation of both mRNA and protein levels of AQP1 (Fig 5A and 7A). These observations could be due to the suppression of RAS by HS diet as Ang II has been reported to increases AQP1 expression in the proximal tubule via direct interaction with angiotensin type 1 receptor [81]. While, all the RAS components are expressed in renal proximal tubule cells [82] suppression of the function of RAS by HS diet may lead to low production of Ang II [83], which has been associated with the downregulation of AQP1 and AQP2 [79].

Perturbation of RAS might also explain the downregulation of AQP2 in the present study. There is evidence showing the relationship between Ang II and AVP. The Ang II increases the secretion of AVP from posterior pituitary which in turn stimulates V_2_ receptor in inner medullary collecting duct [84-87]. In addition, AQP2 is well recognised as AVP-regulated water channel that is expressed in the principal cell of collecting duct. It plays a key role in urine concentration and body-water homeostasis through short- and long-term regulations of water permeability at the collecting duct [88-92]. The AVP through a cascade of events leads to trafficking and marked increased level of AQP2 via gene transcription as well as protein degradation on basolateral membrane. This leads to an increase in permeability to water [28, 31, 93, 94]. The low AQP2 level in the present study could be due to the compensatory mechanism other than via AVP; though plasma AVP was slightly higher (Fig 3D) in SHRs. Meanwhile, the low mRNA and protein levels of AQP2 in WKY rats as result of HS diet may directly due to the low AVP in these rats. Nevertheless, the observation in WKY rats in the present study is in accordance with the study by Roxas et al. [93], which showed that low expression of AQP2 transcript in SD rats fed with HS diet. In addition, stimulation of thirst by HS diet may also be a possible explanation for the suppressed AQP2 in both strains of rats which excessive water drinking keeps circulating AVP levels very low, resulting presumably in suppressed AQP2 levels in the kidneys [95].

Both AQP3 and AQP4 are constitutively localised in basolateral membrane in principal cells of collecting duct. To be more precise, AQP3 is found in cortical and outer medullary collecting duct, whereas AQP4 is located primarily in inner medullary collecting ducts. They both represent potential exit pathways i.e. the increased intracellular water absorbed by AQP2 is transported to blood by AQP3 and AQP4 [35] according to an osmotic gradient. In the present study, both these AQPs showed upregulation in mRNA expression level in both strains of rats fed with HS diet (Fig 5C and D); whilst, SHRs showed lower protein expression of AQP4. The dramatic upregulation of AQP3 and AQP4 mRNA expression as a consequence of HS diet indicates that the increased water reabsorption in collecting duct may contribute to extracellular volume expansion, which is a typical characteristic of SSH. This is further supported by our findings (Fig 1) that showed the higher MAP in SHRs and WKY rats consuming HS diet. Furthermore, SHRs are known to have a high AQP3 level [27, 32]. The upregulation of AQP3 is in consistent with higher protein expression of AQP3 in the present study (Fig 7C). However, the downregulation of protein expression of AQP4 in SHRs remains to be elucidated.

On the other hand, AQP7 localised at the brush border of proximal straight tubule where AQP1 is also located has been classified as aquaglyceroporins because of its credibility to transports water and glycerol as well as urea just as AQP3. In the present study, expressions of AQP7 at mRNA and protein level (Fig 5F and 7F) were low in both strains of rats being fed with HS diet. The changes in mRNA and protein expressions of AQP7 are in a similar manner as that of AQP1 suggesting a substantial contribution of AQP7 in water reabsorption in the proximal tubule. This observation is in supportive with the study by Sohara et al [96] that showed *AQP-1/AQP-7* double knockout mice showed reduced urinary concentrating ability compared with *AQP-1* solo knockout mice. However, compared to AQP1 the contribution of AQP7 to water permeability in proximal tubule is small and remains to be further examined.

Meanwhile, the AQP6 which has been known to have low water permeability, acting mainly as an anion transporter, is thought to be involved in urinary acid secretion [5, 6, 97]. Furthermore, AQP6 is co-localised with H+ ATPase, suggesting that low pH could activate the protein. These indicate that AQP6 is most likely not involved in transepithelial water transport [98]; therefore, the vice versa regulation in mRNA (Fig 5E and 7E) levels of AQP6 as a consequence of HS diet hugely remains unexplained.

## Concluding Remarks

In summary, HS diet intake markedly increased MAP in SHRs and this increase does not seem to be associated with renal expressions of ENaC and AQP subunits. The lower expression and distribution of ENaC and AQP subunits as a consequence of HS intake suggest stimulation of BP regulatory system in SHRs in an attempt to maintain the MAP; and here it is likely via natriuresis activated by ANP. A significant higher plasma ANP activity and lower plasma aldosterone level seen in the present study strongly correlate with the suppression of ENaC and AQP subunits. Furthermore, the present finding suggests that the kidney sodium- and water-handling channels may not directly responsible for the increase in MAP by HS diet intake in SHRs. Thus, the role of ENaC and AQP subunits in salt-sensitive HPN is more towards the maintenance of BP rather than rising the BP.

## Authors’ Contribution Statement

S-ZH, KG, MRM and S-KL conceived the study and assisted in manuscript editing. CDR conducted the experiments and wrote the initial draft of the manuscript.

## Acknowledgements

University Malaya and Ministry of Higher Education, Malaysia

## References

1. Johnson AK, Zhang Z, Clayton SC, Beltz TG, Hurley SW, Thunhorst RL, et al. The roles of sensitization and neuroplasticity in the long-term regulation of blood pressure and hypertension. Am J Physiol Regul Integr Comp Physiol. 2015;309(11):R1309–25. doi: 10.1152/ajpregu.00037.2015. PubMed PMID: 26290101; PubMed Central PMCID: PMCPMC4698407.

2. Blaustein MP, Leenen FH, Chen L, Golovina VA, Hamlyn JM, Pallone TL, et al. How NaCl raises blood pressure: a new paradigm for the pathogenesis of salt-dependent hypertension. American journal of physiology Heart and circulatory physiology. 2012;302(5):H1031–49. doi: 10.1152/ajpheart.00899.2011. PubMed PMID: 22058154; PubMed Central PMCID: PMCPMC3311458.

3. Hall JE. Kidney Dysfunction Mediates Salt-Induced Increases in Blood Pressure. Circulation. 2016;133(9):894–906.

4. Busst CJ. Blood pressure regulation via the epithelial sodium channel: from gene to kidney and beyond. Clin Exp Pharmacol Physiol. 2013;40(8):495–503. doi: 10.1111/1440-1681.12124. PubMed PMID: 23710770.

5. Esteva-Font C, Ballarin J, Fernandez-Llama P. Molecular biology of water and salt regulation in the kidney. Cell Mol Life Sci. 2012;69(5):683–95. doi: 10.1007/s00018-011-0858-4. PubMed PMID: 21997386.

6. Kim HY. Renal Sodium Transporters and Water Channels. J Korean Soc Hypertens. 2013;19(1):17–22.

7. Canessa CM, Schild L, Buell G, Thorens B, Gautschi I, Horisberger JD, et al. Amiloride-sensitive epithelial Na^+^ channel is made of three homologous subunits. Nature. 1994;367(6462):463–7. doi: 10.1038/367463a0. PubMed PMID: 8107805.

8. Schild L. The epithelial sodium channel and the control of sodium balance. Biochim Biophys Acta. 2010;1802(12):1159–65. doi: 10.1016/j.bbadis.2010.06.014. PubMed PMID: 20600867.

9. Bhalla V, Hallows KR. Mechanisms of ENaC regulation and clinical implications. Journal of the American Society of Nephrology: JASN. 2008;19(10):1845–54. doi: 10.1681/ASN.2008020225. PubMed PMID: 18753254.

10. Kleyman TR, Carattino MD, Hughey RP. ENaC at the cutting edge: regulation of epithelial sodium channels by proteases. J Biol Chem. 2009;284(31):20447–51. doi: 10.1074/jbc.R800083200. PubMed PMID: 19401469; PubMed Central PMCID: PMCPMC2742807.

11. Soundararajan R, Pearce D, Hughey RP, Kleyman TR. Role of epithelial sodium channels and their regulators in hypertension. J Biol Chem. 2010;285(40):30363–9. doi: 10.1074/jbc.R110.155341. PubMed PMID: 20624922; PubMed Central PMCID: PMCPMC2945528.

12. Garty H. Regulation of the epithelial Na^+^ channel by aldosterone: open questions and emerging answers. Kidney Int. 2000;57(4):1270–6. doi: 10.1046/j.1523-1755.2000.00961.x. PubMed PMID: 10760053.

13. Garty H, Palmer LG. Epithelial sodium channels: function, structure, and regulation. Physiol Rev. 1997;77(2):359–96. doi: 10.1152/physrev.1997.77.2.359. PubMed PMID: 9114818.

14. Bankir L, Bichet DG, Bouby N. Vasopressin V2 receptors, ENaC, and sodium reabsorption: a risk factor for hypertension? American journal of physiology Renal physiology. 2010;299(5):F917–28. doi: 10.1152/ajprenal.00413.2010. PubMed PMID: 20826569.

15. Reif MC, Troutman SL, Schafer JA. Sustained response to vasopressin in isolated rat cortical collecting tubule. Kidney Int. 1984;26(5):725–32. PubMed PMID: 6097738.

16. Tomita K, Pisano JJ, Knepper MA. Control of sodium and potassium transport in the cortical collecting duct of the rat. Effects of bradykinin, vasopressin, and deoxycorticosterone. J Clin Invest. 1985;76(1):132–6. doi: 10.1172/JCI111935. PubMed PMID: 4019771; PubMed Central PMCID: PMCPMC423727.

17. Beutler KT, Masilamani S, Turban S, Nielsen J, Brooks HL, Ageloff S, et al. Long-term regulation of ENaC expression in kidney by angiotensin II. Hypertension. 2003;41(5):1143–50. doi: 10.1161/01.HYP.0000066129.12106.E2. PubMed PMID: 12682079.

18. Peti-Peterdi J, Warnock DG, Bell PD. Angiotensin II directly stimulates ENaC activity in the cortical collecting duct via AT(1) receptors. Journal of the American Society of Nephrology: JASN. 2002;13(5):1131–5. PubMed PMID: 11960999.

19. Wang Q, Horisberger JD, Maillard M, Brunner HR, Rossier BC, Burnier M. Salt-and angiotensin II-dependent variations in amiloride-sensitive rectal potential difference in mice. Clin Exp Pharmacol Physiol. 2000;27(1-2):60–6. PubMed PMID: 10696530.

20. Guo LJ, Alli AA, Eaton DC, Bao HF. ENaC is regulated by natriuretic peptide receptor-dependent cGMP signaling. American journal of physiology Renal physiology. 2013;304(7):F930–7. doi: 10.1152/ajprenal.00638.2012. PubMed PMID: 23324181; PubMed Central PMCID: PMCPMC4073950.

21. Sun Y, Zhang JN, Zhao D, Wang QS, Gu YC, Ma HP, et al. Role of the epithelial sodium channel in salt-sensitive hypertension. Acta Pharmacol Sin. 2011;32(6):789–97. doi: 10.1038/aps.2011.72. PubMed PMID: 21623391; PubMed Central PMCID: PMCPMC4009973.

22. Aoi W, Niisato N, Sawabe Y, Miyazaki H, Tokuda S, Nishio K, et al. Abnormal expression of ENaC and SGK1 mRNA induced by dietary sodium in Dahl salt-sensitively hypertensive rats. Cell Biol Int. 2007;31(10):1288–91. doi: 10.1016/j.cellbi.2007.03.036. PubMed PMID: 17485228.

23. Dahl LK, Heine M, Tassinari L. Effects of chronia excess salt ingestion. Evidence that genetic factors play an important role in susceptibility to experimental hypertension. J Exp Med. 1962;115:1173–90. PubMed PMID: 13883089; PubMed Central PMCID: PMCPMC2137393.

24. Fenton RA, Chou CL, Ageloff S, Brandt W, Stokes JB, Knepper MA. Increased collecting duct urea transporter expression in Dahl salt-sensitive rats. American journal of physiology Renal physiology. 2003;285(1):F143–51. doi: 10.1152/ajprenal.00073.2003. PubMed PMID: 12684228.

25. Preston GM, Carroll TP, Guggino WB, Agre P. Appearance of water channels in Xenopus oocytes expressing red cell CHIP28 protein. Science. 1992;256(5055):385–7. PubMed PMID: 1373524.

26. Agarwal SK, Gupta A. Aquaporins: The renal water channels. Indian J Nephrol. 2008;18(3):95–100. doi: 10.4103/0971-4065.43687. PubMed PMID: 20142913; PubMed Central PMCID: PMCPMC2813137.

27. Procino G, Romano F, Torielli L, Ferrari P, Bianchi G, Svelto M, et al. Altered expression of renal aquaporins and alpha-adducin polymorphisms may contribute to the establishment of salt-sensitive hypertension. American journal of hypertension. 2011;24(7):822–8. doi: 10.1038/ajh.2011.47. PubMed PMID: 21451595.

28. Buemi M, Nostro L, Di Pasquale G, Cavallaro E, Sturiale A, Floccari F, et al. Aquaporin-2 water channels in spontaneously hypertensive rats. American journal of hypertension. 2004;17(12 Pt 1):1170–8. doi: 10.1016/j.amjhyper.2004.07.003. PubMed PMID: 15607625.

29. Nejsum LN, Elkjaer M, Hager H, Frokiaer J, Kwon TH, Nielsen S. Localization of aquaporin-7 in rat and mouse kidney using RT-PCR, immunoblotting, and immunocytochemistry. Biochem Biophys Res Commun. 2000;277(1):164–70. doi: 10.1006/bbrc.2000.3638. PubMed PMID: 11027658.

30. Nielsen S, Frokiaer J, Marples D, Kwon TH, Agre P, Knepper MA. Aquaporins in the kidney: from molecules to medicine. Physiol Rev. 2002;82(1):205–44. doi: 10.1152/physrev.00024.2001. PubMed PMID: 11773613.

31. Graffe CC, Bech JN, Pedersen EB. Effect of high and low sodium intake on urinary aquaporin-2 excretion in healthy humans. American journal of physiology Renal physiology. 2012;302(2):F264–75. doi: 10.1152/ajprenal.00442.2010. PubMed PMID: 21993890.

32. Lee J, Kim S, Kim J, Jeong MH, Oh Y, Choi KC. Increased expression of renal aquaporin water channels in spontaneously hypertensive rats. Kidney Blood Press Res. 2006;29(1):18–23. doi: 10.1159/000092483. PubMed PMID: 16582573.

33. Kortenoeven ML, Fenton RA. Renal aquaporins and water balance disorders. Biochim Biophys Acta. 2014;1840(5):1533–49. doi: 10.1016/j.bbagen.2013.12.002. PubMed PMID: 24342488.

34. Matsuzaki T, Yaguchi T, Shimizu K, Kita A, Ishibashi K, Takata K. The distribution and function of aquaporins in the kidney: resolved and unresolved questions. Anat Sci Int. 2017;92(2):187–99. doi: 10.1007/s12565-016-0325-2. PubMed PMID: 26798062.

35. Kim JM, Kim TH, Wang T. Effect of Diet and Water Intake on Aquaporin 2 Function. Child Kidney Dis. 2016;20:11–7.

36. Gutkowska J, Horky K, Thibault G, Januszewicz P, Cantin M, Genest J. Atrial natriuretic factor is a circulating hormone. Biochem Biophys Res Commun. 1984;125(1):315–23. PubMed PMID: 6542365.

37. Livak KJ, Schmittgen TD. Analysis of relative gene expression data using real-time quantitative PCR and the 2(-Delta Delta C(T)) Method. Methods. 2001;25(4):402–8. doi: 10.1006/meth.2001.1262. PubMed PMID: 11846609.

38. Lee SW, Schwartz A, Adams RJ, Yamori Y, Whitmer K, Lane LK, et al. Decrease in Na^+^,K+-ATPase activity and [3H]ouabain binding sites in sarcolemma prepared from hearts of spontaneously hypertensive rats. Hypertension. 1983;5(5):682–8. PubMed PMID: 6311739.

39. Sato Y, Ando K, Ogata E, Fujita T. Salt sensitivity in Goldblatt hypertensive rats--role of extracellular fluid volume and renin-angiotensin system. Japanese circulation journal. 1991;55(2):165–73. PubMed PMID: 2020087.

40. de Wardener HE, He FJ, MacGregor GA. Plasma sodium and hypertension. Kidney Int. 2004;66(6):2454–66. doi: 10.1111/j.1523-1755.2004.66018.x. PubMed PMID: 15569339.

41. Fitzsimons JT. Angiotensin, thirst, and sodium appetite. Physiol Rev. 1998;78(3):583–686. PubMed PMID: 9674690.

42. Qi N, Rapp JP, Brand PH, Metting PJ, Britton SL. Body fluid expansion is not essential for salt-induced hypertension in SS/Jr rats. Am J Physiol. 1999;277(5 Pt 2):R1392–400. PubMed PMID: 10564212.

43. De Luca LA, Jr., Pereira-Derderian DT, Vendramini RC, David RB, Menani JV. Water deprivation-induced sodium appetite. Physiol Behav. 2010;100(5):535–44. doi: 10.1016/j.physbeh.2010.02.028. PubMed PMID: 20226201.

44. Wolf K, Kurtz A. Influence of salt intake on atrial natriuretic peptide gene expression in rats. Pflugers Arch. 1997;433(6):809–16. doi: 10.1007/s004240050349. PubMed PMID: 9049174.

45. Greenwood MP, Mecawi AS, Hoe SZ, Mustafa MR, Johnson KR, Al-Mahmoud GA, et al. A comparison of physiological and transcriptome responses to water deprivation and salt loading in the rat supraoptic nucleus. Am J Physiol Regul Integr Comp Physiol. 2015;308(7):R559–68. doi: 10.1152/ajpregu.00444.2014. PubMed PMID: 25632023; PubMed Central PMCID: PMCPMC4386000.

46. Sagnella GA, Markandu ND, Buckley MG, Miller MA, Singer DR, Cappuccio FP, et al. Atrial natriuretic peptides in essential hypertension: basal plasma levels and relationship to sodium balance. Can J Physiol Pharmacol. 1991;69(10):1592–600. PubMed PMID: 1838027.

47. Kohno M, Yasunari K, Murakawa K, Kanayama Y, Matsuura T, Takeda T. Effects of high-sodium and low-sodium intake on circulating atrial natriuretic peptides in salt-sensitive patients with systemic hypertension. Am J Cardiol. 1987;59(12):1212–3. PubMed PMID: 2953231.

48. Antunes-Rodrigues J, de Castro M, Elias LL, Valenca MM, McCann SM. Neuroendocrine control of body fluid metabolism. Physiol Rev. 2004;84(1):169–208. doi: 10.1152/physrev.00017.2003. PubMed PMID: 14715914.

49. Candela L, Yucha C. Renal regulation of extracellular fluid volume and osmolality. Nephrol Nurs J. 2004;31(4):397–404, 44; quiz 5-6. PubMed PMID: 15453232.

50. Kaschina E, Unger T. Angiotensin AT1/AT2 receptors: regulation, signalling and function. Blood Press. 2003;12(2):70–88. PubMed PMID: 12797627.

51. Mecawi AS, Vilhena-Franco T, Fonseca FV, Reis LC, Elias LL, Antunes-Rodrigues J. The role of angiotensin II on sodium appetite after a low-sodium diet. J Neuroendocrinol. 2013;25(3):281–91. doi: 10.1111/j.1365-2826.2012.02388.x. PubMed PMID: 23002791.

52. Oki K, Gomez-Sanchez EP, Gomez-Sanchez CE. Role of mineralocorticoid action in the brain in salt-sensitive hypertension. Clin Exp Pharmacol Physiol. 2012;39(1):90–5. doi: 10.1111/j.1440-1681.2011.05538.x. PubMed PMID: 21585422; PubMed Central PMCID: PMCPMC3164934.

53. Cowley AW, Jr., Cushman WC, Quillen EW, Jr., Skelton MM, Langford HG. Vasopressin elevation in essential hypertension and increased responsiveness to sodium intake. Hypertension. 1981;3(3 Pt 2):I93–100. PubMed PMID: 7262983.

54. Mohring J, Kintz J, Schoun J. Studies on the role of vasopressin in blood pressure control of spontaneously hypertensive rats with established hypertension (SHR, stroke-prone strain). J Cardiovasc Pharmacol. 1979;1(6):593–608. PubMed PMID: 94626.

55. Kawano Y, Matsuoka H, Nishikimi T, Takishita S, Omae T. The role of vasopressin in essential hypertension. Plasma levels and effects of the V1 receptor antagonist OPC-21268 during different dietary sodium intakes. American journal of hypertension. 1997;10(11):1240–4. PubMed PMID: 9397242.

56. Dilley JR, Stier CT, Jr., Arendshorst WJ. Abnormalities in glomerular function in rats developing spontaneous hypertension. Am J Physiol. 1984;246(1 Pt 2):F12–20. doi: 10.1152/ajprenal.1984.246.1.F12. PubMed PMID: 6696074.

57. Kim SW, Wang W, Kwon TH, Knepper MA, Frokiaer J, Nielsen S. Increased expression of ENaC subunits and increased apical targeting of AQP2 in the kidneys of spontaneously hypertensive rats. American journal of physiology Renal physiology. 2005;289(5):F957–68. doi: 10.1152/ajprenal.00413.2004. PubMed PMID: 15956775.

58. Tahara A, Tsukada J, Tomura Y, Wada K, Kusayama T, Ishii N, et al. Alterations of renal vasopressin V1A and V2 receptors in spontaneously hypertensive rats. Pharmacology. 2003;67(2):106–12. doi: 10.1159/000067743. PubMed PMID: 12566855.

59. Morduchowicz GA, Sheikh-Hamad D, Jo OD, Nord EP, Lee DB, Yanagawa N. Increased Na^+^/H+ antiport activity in the renal brush border membrane of SHR. Kidney Int. 1989;36(4):576–81. PubMed PMID: 2554051.

60. Naray-Fejes-Toth A, Canessa C, Cleaveland ES, Aldrich G, Fejes-Toth G. sgk is an aldosterone-induced kinase in the renal collecting duct. Effects on epithelial Na^+^ channels. J Biol Chem. 1999;274(24):16973–8. PubMed PMID: 10358046.

61. Palmer LG. Regulation of epithelial Na channels by aldosterone. Kitasato Med J. 2016;46:1–7.

62. Verrey F, Fakitsas P, Adam G, Staub O. Early transcriptional control of ENaC (de)ubiquitylation by aldosterone. Kidney Int. 2008;73(6):691–6. doi: 10.1038/sj.ki.5002737. PubMed PMID: 18094676.

63. Escoubet B, Coureau C, Bonvalet JP, Farman N. Noncoordinate regulation of epithelial Na channel and Na pump subunit mRNAs in kidney and colon by aldosterone. Am J Physiol. 1997;272(5 Pt 1):C1482–91. doi: 10.1152/ajpcell.1997.272.5.C1482. PubMed PMID: 9176138.

64. Masilamani S, Wang X, Kim GH, Brooks H, Nielsen J, Nielsen S, et al. Time course of renal Na-K-ATPase, NHE3, NKCC2, NCC, and ENaC abundance changes with dietary NaCl restriction. American journal of physiology Renal physiology. 2002;283(4):F648–57. doi: 10.1152/ajprenal.00016.2002. PubMed PMID: 12217855.

65. Shehata MF. Regulation of the epithelial sodium channel [ENaC] in kidneys of salt-sensitive Dahl rats: insights on alternative splicing. Int Arch Med. 2009;2(1):28. doi: 10.1186/1755-7682-2-28. PubMed PMID: 19785774; PubMed Central PMCID: PMCPMC2761857.

66. Hamm LL, Feng Z, Hering-Smith KS. Regulation of sodium transport by ENaC in the kidney. Current opinion in nephrology and hypertension. 2010;19(1):98–105. doi: 10.1097/MNH.0b013e328332bda4. PubMed PMID: 19996890; PubMed Central PMCID: PMCPMC2895494.

67. Barker PM, Nguyen MS, Gatzy JT, Grubb B, Norman H, Hummler E, et al. Role of gammaENaC subunit in lung liquid clearance and electrolyte balance in newborn mice. Insights into perinatal adaptation and pseudohypoaldosteronism. J Clin Invest. 1998;102(8):1634–40. doi: 10.1172/JCI3971. PubMed PMID: 9788978; PubMed Central PMCID: PMCPMC509015.

68. Huang BS, Van Vliet BN, Leenen FH. Increases in CSF [Na^+^] precede the increases in blood pressure in Dahl S rats and SHR on a high-salt diet. American journal of physiology Heart and circulatory physiology. 2004;287(3):H1160–6. doi: 10.1152/ajpheart.00126.2004. PubMed PMID: 15130889.

69. Nakano M, Hirooka Y, Matsukawa R, Ito K, Sunagawa K. Mineralocorticoid receptors/epithelial Na(+) channels in the choroid plexus are involved in hypertensive mechanisms in stroke-prone spontaneously hypertensive rats. Hypertens Res. 2013;36(3):277–84. doi: 10.1038/hr.2012.174. PubMed PMID: 23096235.

70. Ritz E, Mehls O. Salt restriction in kidney disease--a missed therapeutic opportunity? Pediatr Nephrol. 2009;24(1):9–17. doi: 10.1007/s00467-008-0856-4. PubMed PMID: 18535843; PubMed Central PMCID: PMCPMC2644745.

71. Farjah M, Roxas BP, Geenen DL, Danziger RS. Dietary salt regulates renal SGK1 abundance: relevance to salt sensitivity in the Dahl rat. Hypertension. 2003;41(4):874–8. doi: 10.1161/01.HYP.0000063885.48344.EA. PubMed PMID: 12642512.

72. Kakizoe Y, Kitamura K, Ko T, Wakida N, Maekawa A, Miyoshi T, et al. Aberrant ENaC activation in Dahl salt-sensitive rats. Journal of hypertension. 2009;27(8):1679–89. doi: 10.1097/HJH.0b013e32832c7d23. PubMed PMID: 19458538.

73. Frindt G, Ergonul Z, Palmer LG. Surface expression of epithelial Na channel protein in rat kidney. J Gen Physiol. 2008;131(6):617–27. doi: 10.1085/jgp.200809989. PubMed PMID: 18504317; PubMed Central PMCID: PMCPMC2391254.

74. Loffing J, Pietri L, Aregger F, Bloch-Faure M, Ziegler U, Meneton P, et al. Differential subcellular localization of ENaC subunits in mouse kidney in response to high- and low-Na diets. American journal of physiology Renal physiology. 2000;279(2):F252–8. doi: 10.1152/ajprenal.2000.279.2.F252. PubMed PMID: 10919843.

75. Chang SY, Lo CS, Zhao XP, Liao MC, Chenier I, Bouley R, et al. Overexpression of angiotensinogen downregulates aquaporin 1 expression via modulation of Nrf2-HO-1 pathway in renal proximal tubular cells of transgenic mice. Journal of the renin-angiotensin-aldosterone system: JRAAS. 2016;17(3). doi: 10.1177/1470320316668737. PubMed PMID: 27638854; PubMed Central PMCID: PMCPMC5843896.

76. Noda M. The subfornical organ, a specialized sodium channel, and the sensing of sodium levels in the brain. Neuroscientist. 2006;12(1):80–91. doi: 10.1177/1073858405279683. PubMed PMID: 16394195.

77. Liu C, Song Y, Qu L, Tang J, Meng L, Wang Y. Involvement of NOX in the regulation of renal tubular expression of Na/K-ATPase in acute unilateral ureteral obstruction rats. Nephron. 2015;130(1):66–76. doi: 10.1159/000381858. PubMed PMID: 25997532.

78. Jiang Y, Wang HY, Zheng S, Mu SQ, Ma MN, Xie X, et al. Cardioprotective effect of valsartan in mice with short-term high-salt diet by regulating cardiac aquaporin 1 and angiogenic factor expression. Cardiovasc Pathol. 2015;24(4):224–9. doi: 10.1016/j.carpath.2014.12.003. PubMed PMID: 25659450.

79. Della Penna SL, Cao G, Fellet A, Balaszczuk AM, Zotta E, Cerrudo C, et al. Salt-induced downregulation of renal aquaporins is prevented by losartan. Regul Pept. 2012;177(1-3):85–91. doi: 10.1016/j.regpep.2012.05.090. PubMed PMID: 22587908.

80. Tomassoni D, Bramanti V, Amenta F. Expression of aquaporins 1 and 4 in the brain of spontaneously hypertensive rats. Brain Res. 2010;1325:155–63. doi: 10.1016/j.brainres.2010.02.023. PubMed PMID: 20156423.

81. Bouley R, Palomino Z, Tang SS, Nunes P, Kobori H, Lu HA, et al. Angiotensin II and hypertonicity modulate proximal tubular aquaporin 1 expression. American journal of physiology Renal physiology. 2009;297(6):F1575–86. doi: 10.1152/ajprenal.90762.2008. PubMed PMID: 19776169; PubMed Central PMCID: PMCPMC2801332.

82. Tang SS, Jung F, Diamant D, Brown D, Bachinsky D, Hellman P, et al. Temperature-sensitive SV40 immortalized rat proximal tubule cell line has functional renin-angiotensin system. Am J Physiol. 1995;268(3 Pt 2):F435–46. doi: 10.1152/ajprenal.1995.268.3.F435. PubMed PMID: 7900843.

83. Drenjancevic-Peric I, Jelakovic B, Lombard JH, Kunert MP, Kibel A, Gros M. High-salt diet and hypertension: focus on the renin-angiotensin system. Kidney Blood Press Res. 2011;34(1):1–11. doi: 10.1159/000320387. PubMed PMID: 21071956; PubMed Central PMCID: PMCPMC3214830.

84. Lee YJ, Song IK, Jang KJ, Nielsen J, Frokiaer J, Nielsen S, et al. Increased AQP2 targeting in primary cultured IMCD cells in response to angiotensin II through AT1 receptor. American journal of physiology Renal physiology. 2007;292(1):F340–50. doi: 10.1152/ajprenal.00090.2006. PubMed PMID: 16896188.

85. Li C, Wang W, Rivard CJ, Lanaspa MA, Summer S, Schrier RW. Molecular mechanisms of angiotensin II stimulation on aquaporin-2 expression and trafficking. American journal of physiology Renal physiology. 2011;300(5):F1255–61. doi: 10.1152/ajprenal.00469.2010. PubMed PMID: 21325494; PubMed Central PMCID: PMCPMC3094043.

86. Torp M, Brond L, Hadrup N, Nielsen JB, Praetorius J, Nielsen S, et al. Losartan decreases vasopressin-mediated cAMP accumulation in the thick ascending limb of the loop of Henle in rats with congestive heart failure. Acta physiologica. 2007;190(4):339–50. doi: 10.1111/j.1748-1716.2007.01722.x. PubMed PMID: 17635349.

87. Wong NL, Tsui JK. Angiotensin II upregulates the expression of vasopressin V2 mRNA in the inner medullary collecting duct of the rat. Metabolism. 2003;52(3):290–5. doi: 10.1053/meta.2003.50047. PubMed PMID: 12647265.

88. DiGiovanni SR, Nielsen S, Christensen EI, Knepper MA. Regulation of collecting duct water channel expression by vasopressin in Brattleboro rat. Proc Natl Acad Sci U S A. 1994;91(19):8984–8. PubMed PMID: 7522327; PubMed Central PMCID: PMCPMC44731.

89. Ecelbarger CA, Terris J, Frindt G, Echevarria M, Marples D, Nielsen S, et al. Aquaporin-3 water channel localization and regulation in rat kidney. Am J Physiol. 1995;269(5 Pt 2):F663–72. doi: 10.1152/ajprenal.1995.269.5.F663. PubMed PMID: 7503232.

90. Nielsen S, Chou CL, Marples D, Christensen EI, Kishore BK, Knepper MA. Vasopressin increases water permeability of kidney collecting duct by inducing translocation of aquaporin-CD water channels to plasma membrane. Proc Natl Acad Sci U S A. 1995;92(4):1013–7. PubMed PMID: 7532304; PubMed Central PMCID: PMCPMC42627.

91. Terris J, Ecelbarger CA, Nielsen S, Knepper MA. Long-term regulation of four renal aquaporins in rats. Am J Physiol. 1996;271(2 Pt 2):F414–22. doi: 10.1152/ajprenal.1996.271.2.F414. PubMed PMID: 8770174.

92. Yamamoto T, Sasaki S, Fushimi K, Ishibashi K, Yaoita E, Kawasaki K, et al. Vasopressin increases AQP-CD water channel in apical membrane of collecting duct cells in Brattleboro rats. Am J Physiol. 1995;268(6 Pt 1):C1546–51. doi: 10.1152/ajpcell.1995.268.6.C1546. PubMed PMID: 7541941.

93. Roxas B, Farjah M, Danziger RS. Aquaporin-2 transcript is differentially regulated by dietary salt in Sprague-Dawley and Dahl SS/Jr rats. Biochem Biophys Res Commun. 2002;296(3):755–8. PubMed PMID: 12176047.

94. Song J, Hu X, Shi M, Knepper MA, Ecelbarger CA. Effects of dietary fat, NaCl, and fructose on renal sodium and water transporter abundances and systemic blood pressure. American journal of physiology Renal physiology. 2004;287(6):F1204–12. doi: 10.1152/ajprenal.00063.2004. PubMed PMID: 15304371.

95. Radin MJ, Yu MJ, Stoedkilde L, Miller RL, Hoffert JD, Frokiaer J, et al. Aquaporin-2 regulation in health and disease. Vet Clin Pathol. 2012;41(4):455–70. doi: 10.1111/j.1939-165x.2012.00488.x. PubMed PMID: 23130944; PubMed Central PMCID: PMCPMC3562700.

96. Sohara E, Rai T, Sasaki S, Uchida S. Physiological roles of AQP7 in the kidney: Lessons from AQP7 knockout mice. Biochim Biophys Acta. 2006;1758(8):1106–10. doi: 10.1016/j.bbamem.2006.04.002. PubMed PMID: 16860289.

97. Verkman AS. Aquaporins: translating bench research to human disease. J Exp Biol. 2009;212(Pt 11):1707–15. doi: 10.1242/jeb.024125. PubMed PMID: 19448080; PubMed Central PMCID: PMCPMC2683014.

98. Yasui M, Kwon TH, Knepper MA, Nielsen S, Agre P. Aquaporin-6: An intracellular vesicle water channel protein in renal epithelia. Proc Natl Acad Sci U S A. 1999;96(10):5808–13. PubMed PMID: 10318966; PubMed Central PMCID: PMCPMC21942.

